# ReCodLiver0.9: Overcoming challenges in genome-scale metabolic reconstruction of a non-model species

**DOI:** 10.1101/2020.06.23.162792

**Authors:** Eileen Marie Hanna, Xiaokang Zhang, Marta Eide, Shirin Fallahi, Tomasz Furmanek, Fekadu Yadetie, Daniel Craig Zielinski, Anders Goksøyr, Inge Jonassen

## Abstract

The availability of genome sequences, annotations and knowledge of the biochemistry underlying metabolic transformations has led to the generation of metabolic network reconstructions for a wide range of organisms in bacteria, archaea, and eukaryotes. When modeled using mathematical representations, a reconstruction can simulate underlying genotype-phenotype relationships. Accordingly, genome-scale models (GEMs) can be used to predict the response of organisms to genetic and environmental variations. A bottom-up reconstruction procedure typically starts by generating a draft model from existing annotation data on a target organism. For model species, this part of the process can be straightforward, due to the abundant organism-specific biochemical data. However, the process becomes complicated for non-model less-annotated species. In this paper, we present a draft liver reconstruction, ReCodLiver0.9, of Atlantic cod (*Gadus morhua*), a non-model teleost fish, as a practicable guide for cases with comparably few resources. Although the reconstruction is considered a draft version, we show that it already has utility in elucidating metabolic response mechanisms to environmental toxicants by mapping gene expression data of exposure experiments to the resulting model.

**Author summary:** Genome-scale metabolic models (GEMs) are constructed based upon reconstructed networks that are carried out by an organism. The underlying biochemical knowledge in such networks can be transformed into mathematical models that could serve as a platform to answer biological questions. The availability of high-throughput biological data, including genomics, proteomics, and metabolomics data, supports the generation of such models for a large number of organisms. Nevertheless, challenges arise for non-model species which are typically less annotated. In this paper, we discuss these challenges and possible solutions in the context of generation of a draft liver reconstruction of Atlantic cod (*Gadus morhua*). We also show how experimental data, here gene expression data, can be mapped to the resulting model to understand the metabolic response of cod liver to environmental toxicants.

## Introduction

Induced by the availability of next-generation DNA sequencing as well as high-throughput genomic, proteomic, and metabolomic data, genome-scale metabolic models (GEMs) have become fundamental tools in the systems biology of metabolism. To develop a GEM, metabolic genes and their product enzymes are assembled into a network of metabolic reactions carried out by an organism [42]. Such networks can be used as a platform to answer relevant biological questions, by transforming biochemical knowledge into a mathematical format and computing appropriate physiological states [41, 46]. Using Boolean logic representation, gene-protein-reaction (GPR) associations depict direct connections between genotype and metabolic capability [39]. GEMs have a wide scope of applications, among which are the contextualization of omics data, hypothesis-driven discovery, support for metabolic engineering, modeling interactions among cells and organisms, drug targeting, and prediction of enzyme functions [20, 40].

A comprehensive bottom-up protocol to generate high-quality reconstructions was presented a decade ago [50]. It consists of four main phases presented as an exhaustive list of steps. First, a draft reconstruction is generated based on the genome annotation of a target organism and available information in biochemical databases, usually in an automated manner. Noticeably, the quality of the draft model is highly impacted by the quality of the genome annotation, in addition to the abundance of data on the organism in public repositories. The draft is then subject to manual refinement that revises and curates all gene and reaction entries. Next, the reconstruction can be converted into mathematical format which is thereafter analyzed for gap filling and tested for stoichiometric balance. Several computational tools were developed to cover this process with automated functionalities [20]. They primarily vary in programming languages, linked databases, input template models, and supported target organisms.

Marine organisms, including fish, represent important resources for human societies, and will be essential components of a growing blue bioeconomy. As we are entering the UN Decade of Ocean Science for Sustainable Development [45, 54], understanding and predicting the effects of environmental stressors, including xenobiotic exposures, on marine species will be one of the main challenges. In this context, metabolic reconstructions can prove very useful. Notwithstanding, when it comes to teleost GEMs, only two models are presented in the literature, MetaFishNet [31] and ZebraGEM [6, 53]. MetaFishNet covers five fish species which makes it challenging to analyze organism-specific data using this model. In addition, subsystems in this GEM are not interconnected or compartmentalized. The first version of the ZebraGEM, a zebrafish GEM, did not contain GPR associations. A very recent version, ZebraGEM 2.0 [53], now includes those associations, with standardized component names, and expanded reactions list.

The Atlantic cod (*Gadus morhua*) is a key species in the North Atlantic marine ecosystem. Its genome was first published in 2011 [48] and has been followed up by more extensive genome sequencing, enabling novel genomic and transcriptomic assemblies [49] as well as a Trinity assembly (We are submitting the assembly to public repositories, and the link or accession number will come afterwards) and a very recent assembly gadMor3.0 (GenBank assembly accession: GCA 902167405). These resources allow extensive gene mapping and bioinformatic analysis, such as the identification of the gene complement of the Atlantic cod cytochrome P450 superfamily [26] or the total chemical defensome [60]. Due to its habitat near offshore installations as well as coastal industries and communities, the Atlantic cod serves as a relevant model species for studying the effects of and responses to anthropogenic activities [13]. In addition, the lipid-rich liver of cod serves as a magnet for lipophilic environmental pollutants, increasing the bioaccumulation within each fish with age and the biomagnification of compounds through the food chain.

The Atlantic cod has lost the MHC II and related genes of the adaptive immune system [48]. Furthermore, we have recently shown that cod, together with several other teleost fish, have lost the pregnane x receptor (*pxr, nr1i2*) gene from their genomes [15]. This xenosensor is activated by a wide range of drugs and pollutants, and regulates the transcription of genes involved in all phases of the xenobiotic and steroid biotransformation [29]. PXR also mediates regulation of hepatic energy metabolism in humans, by inducing lipogenesis and inhibiting fatty acid beta-oxidation [56]. The absence of a *pxr* gene thus leads to questions regarding how the Atlantic cod responds to environmental contaminants that bind this receptor in other species. Results from an in vivo exposure study, where cod was injected the legacy polychlorinated biphenyl congener PCB153, a compound known to bind PXR in other species, showed indications that lipid metabolism was affected [58]. In view of these findings, an ensuing effort was initiated to generate a metabolic reconstruction of Atlantic cod liver, with a focus on lipid and xenobiotic metabolism pathways. In this paper, we present and discuss our metabolic reconstruction progress in terms of steps, methods, choice of tools, and challenges.

## Methods

### Selecting the appropriate tool

Many computational tools have been developed to help automate steps of the GEM reconstruction process which is usually laborious and time-consuming [50]. We highlight a few approaches. CarveME [34] uses a top-down approach, based on Python command-line, to convert curated reactions from the BiGG database into organism-specific GEMs. AutoKEGGRec [25] is a novel tool which is based on the KEGG pathways database [4] and is fully integrated with the COBRA toolbox [46]. AuReMe [2] presents a customized platform and pipelines to generate GEMs while preserving metadata and ensuring model traceability. It is also linked to the BiGG knowledgebase [14] and the MetaCyc database [11]. For a thorough review of GEM reconstruction tools, we refer to the recent paper by Mendoza et al. [37]. Table 1 lists the tools available for non-model species reconstructions.

**Table 1.**
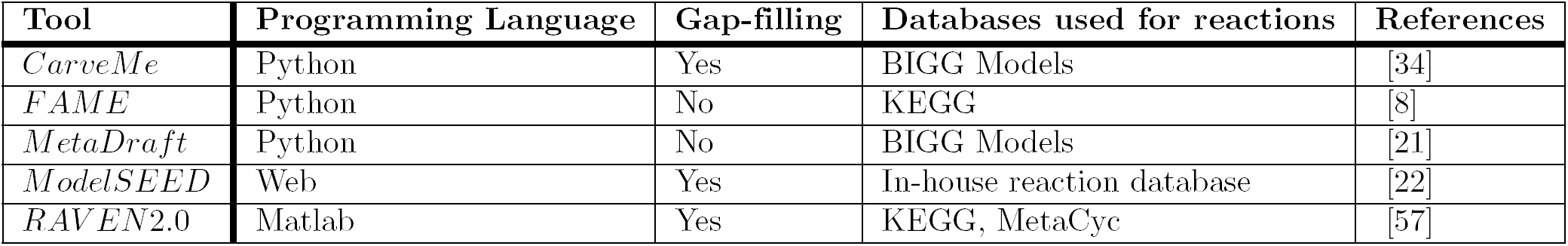
Summary of reconstruction tools for GEMs of non-model species

| Tool | Programming Language | Gap-filling | Databases used for reactions | References |
| --- | --- | --- | --- | --- |
| *CarveMe* | Python | Yes | BIGG Models | [34] |
| *FAME* | Python | No | KEGG | [8] |
| *MetaDraft* | Python | No | BIGG Models | [21] |
| *ModelSEED* | Web | Yes | In-house reaction database | [22] |
| *RAVEN2.0* | Matlab | Yes | KEGG, MetaCyc | [57] |

The choice of tool for our reconstruction work on Atlantic cod was predominantly narrowed down by the fact that this is a non-model and less-annotated organism. De novo reconstruction based on metabolic databases do not apply in such cases, but rather GEMs of related scope needs to serve as input. Another important feature for us was the tool’s support for eukaryote modeling. In view of these constraints, the Reconstruction, Analysis and Visualization of Metabolic Networks (RAVEN 2.0) MATLAB toolbox [57] was selected. It is used for semi-automated draft model reconstruction and provides curation, simulation, and constraint-based capabilities that are compatible with the COBRA toolbox. RAVEN has been used to generate GEMs for several organisms including bacteria, archaea, parasites, fungi, and various human tissues. We also note that the toolbox allows de novo draft reconstruction based on metabolic pathway databases, namely KEGG and MetaCyc, although that it is not applicable to our case study. Most importantly, RAVEN can generate draft models for a target organism based on protein homology, using existing high-quality GEMs of organisms at an appropriate evolutionary distance. Two main functions are applied in this process. getBlast or getBlastFromExcel construct a structure including homology measurements between the target organism and the template organisms. Then, getModelFromHomology uses this structure along with existing template GEMs to create a draft model containing reactions associated with orthologous genes. The subsequent draft is then subject to curation and refinement. The RAVEN toolbox includes several functions that are implemented to support subsequent steps.

### Choosing the reconstruction template

With the Atlantic cod liver as our reconstruction scope, we surveyed the literature for a relevant GEM that could be used as an input template to the RAVEN 2.0 toolbox. Initially, the global reconstruction of the human metabolic network, Recon 1, was published in 2007 [14]. Several network and supporting information updates have followed. A recent Recon 3D version includes a thorough catalogue of GPR associations in addition to structural information on metabolites and enzymes [10]. This version also comprises a three-dimensional metabolite and protein structure data which allow integrated analyses of metabolic functions in humans. The Edinburgh Human Metabolic Network (EHMN) was presented concurrently to Recon 1 [33]. The Human Metabolic Reaction (HMR) database [1] was later generated by integrating Recon 1 and EHMN with biochemical reactions from KEGG [4] and MetaCyc [11] databases. An updated version (HMR 2.0) with emphasis on lipid metabolism was later introduced [36]. Several subsequent tissue- and context-specific GEMs were released. They include iAdipocyte1809, iMyocytes2419 and CardioNet [16, 35]. We emphasize the consensus GEM for hepatocytes, iHepatocyte2322 [36], which was assembled from proteomics data and previously published liver models. It is a consensus hepatocyte GEM that is mainly generated from proteomics data of several existing liver models, namely HepatoNet1 [17], iLJ1046 [24], iAB676 [9], and iHepatocyte1154 [1]. It is well-annotated as it includes all the protein-coding genes and reactions in those liver models. Notably, and in alignment with our GEM context, iHepatocyte2322 extensively covers lipid metabolism functions, such as the uptake of remnants of lipoproteins, the formation and degradation of lipid droplets, and secretion of synthesized lipoproteins, as well as xenobiotic biotransformation. It consists of 2322 genes, 5686 metabolites, and 7930 reactions, separated into eight different compartments based on the HMR 2.0 database. Although the template is not fish-specific, we note that the important subcellular functions and mechanisms within the compartments are assumed conserved through evolution. In contrast to humans that mainly store fat in adipose tissues, Atlantic cod stores most lipids in liver. In fact, cod liver normally contains 50-60% of fat [32] that is stored in peroxisomes and cytosolic lipid droplets, and these subcellular compartments are also included in iHepatocyte2322.

Of the two available fish GEMs, the MetaFishNet [31] involves five fish species: zebrafish (D. rerio), medaka (O. latipes), Takifugu (T. rubripes), Tetraodon (T. nigroviridis) and stickleback (G. aculeatus). Here, cDNA sequences from those species were analyzed and the corresponding list of metabolic genes was formed based on gene ontology. This list was then used to identify enzymes based on their orthology to human genes. Metabolic reactions were integrated from EHMN, the human metabolic network at UCSD (BiGG) [14], and the D. rerio metabolic network from KEGG [4]. The other fish model is ZebraGEM [6]. It is a comprehensive GEM based on the literature and D. rerio biochemical data collected from various resources. This model is manually curated with gaps extensively checked and filled. An updated version, ZebraGEM 2.0, now includes the GPR associations and essential reactions, which were not previously covered in the first version. Both versions simplify the description of lipid metabolism processes by mainly discarding reactions specific to fatty acids. Therefore, essential pathways in our reconstruction scope were excluded from ZebraGEM, which makes it unfit to our case study.

In the absence of tissue-specific models for fish species and the availability of a human liver GEM that greatly focuses on lipid metabolism, a primary trade-off was between using a generic template from a phylogenetically closer species, i.e. zebrafish, or using a liver-specific template from human that falls within our reconstruction scope. The iHepatocyte2322 consensus GEM was our template of choice.

### Generating an Atlantic cod liver draft model

As described, we used the RAVEN 2.0 toolbox to generate a draft metabolic reconstruction of the Atlantic cod liver, based on protein homology, and using iHepatocyte2322 [36] as our input template.

In addition to the template model as input, RAVEN 2.0 requires two FASTA files including the peptide sequences of the target and the template organisms. The literature and public repositories contain four main versions of Atlantic cod genome annotations. The first assembly, gadMor1, was published in 2011 [48]. An updated version, gadMor2, with extensive genome sequencing enabling novel genomic and transcriptomics assemblies followed in 2017 [49]. Those two versions along with multiple RNA-seq data from multiple tissues and developmental stages were used to create gadMorTrinity (reference) which includes less-fragmented and more comprehensive sequences. gadMor3 (GenBank assembly accession: GCA 902167405), here gadMor3, is a recently published chromosome-level reference assembly generated using long-read sequencing technology. Some statistics of the peptide sequences from these three assemblies are listed in Table 2. We used gadMor1, gadMorTrinity, and gadMor3 to examine the potential improvement that these annotations could bring in the context of metabolic reconstructions. A separate set of draft models was generated from each assembly.

**Table 2.** Some statistics of the peptide sequences from three Atlantic cod assemblies

| Statistics | gadMor1 | gadMorTrinity | gadMor3 |
| --- | --- | --- | --- |
| Number of sequences | 22100 | 34737 | 45039 |
| Average length | 480 | 712 | 729 |

First, the FASTA files were used as input to the getBlast function. A BLAST structure, was created from the reciprocal BLAST analyses between the target and the template organism, using the BLASTp algorithm [3]. Next, this structure along with the template model were input to the getModelFromHomology function to generate a draft cod GEM by extracting reactions involving cod genes that are orthologous to human genes in iHepatocyte2322. Fig 1 shows an overview of the draft model reconstruction process. The process can be tuned based on different input parameters.

**Fig 1.**
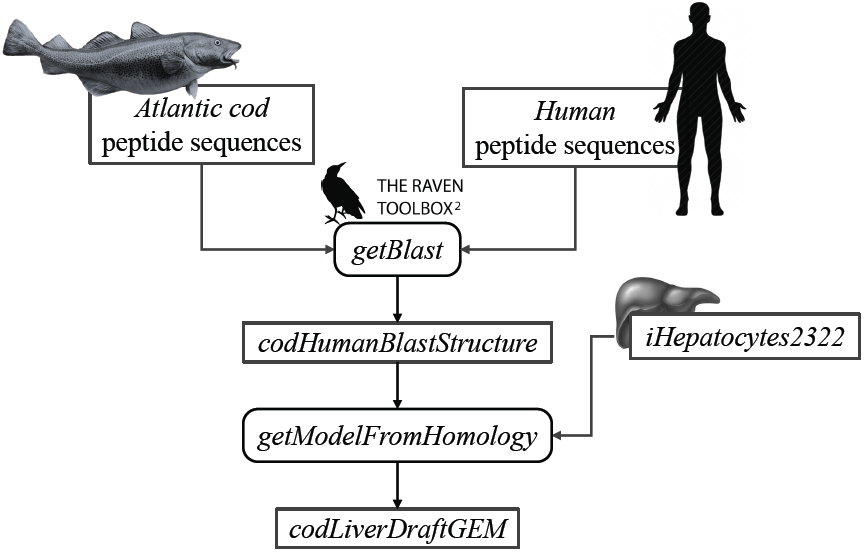
Schematic overview of the reconstruction of an Atlantic cod liver draft model.

We used the RAVEN 2.0 toolbox to generate several draft models based on different strictness parameters and Atlantic cod genome annotations. The strictness parameter specifies which reactions are included in the draft model, mainly based on how genes from the template GEM organism and from the target organism are mapped against each other. When strictness is equal to 1 or 2, gene mapping is done for all pairs based on BLASTp score threshold, in both or in one direction, respectively. Alternatively, when strictness is set to 3, all BLASTp results are checked and only the ones with the lowest E-value are kept for all gene pairs in each direction separately. Then, target genes are mapped to template genes for all pairs which have acceptable BLASTp results, in both directions. In addition, maxE, minLen, and minIde are adjustable thresholds for the maximum E-value, the minimum alignment length, and the minimum identity acceptable in the gene mapping process, respectively.

## Results

The graph in Fig 2 shows the model statistics corresponding to iHepatocytes2322 and different cod annotations, based on strictness values of 1 (s1) and 3 (s3) for one-to-many and best one-to-one mapping between human and cod genes, respectively. First, we note that s1 leads to a greater number of matched genes in the draft models. Such high number of matches can be attributed to several whole-genome duplications during teleost fish evolution, by which fish genomes often have more than one ortholog of human genes [18, 19, 44]. Accordingly, the more annotated the cod genome, the more duplicate matches with human genes in the template are expected. In this context, the trend in the number of genes in the draft models can be attributed to the fact that gadMor1 has the lowest while gadMor3 has the highest number of annotated sequences. The number of reactions and metabolites across all resultant drafts is close for both strictness values.

**Fig 2.**
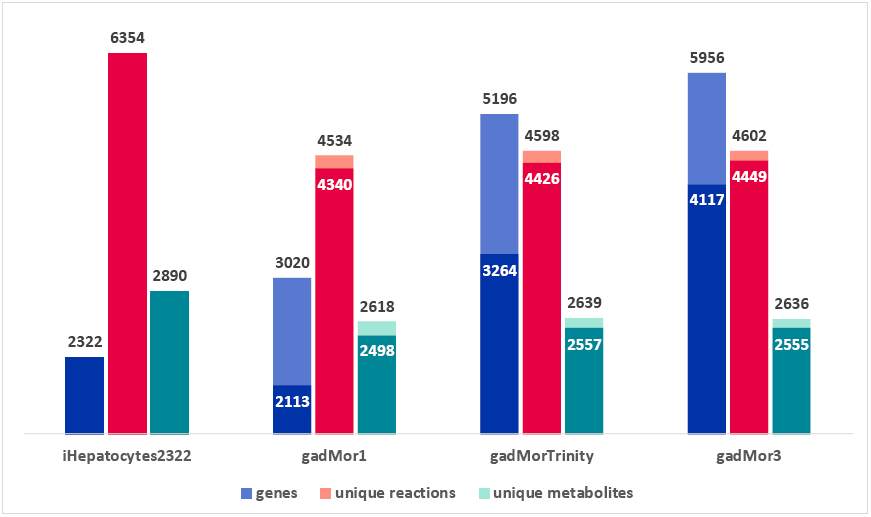
Model statistics of iHepatocytes2322 compared to draft reconstructions of Atlantic cod liver, using different annotations as reference. Dark color indicates strictness of s3 (one to one) and light color indicates strictness of s1 (one to many).

The current version of the RAVEN toolbox adds a reaction to the draft if at least one of its catalyzing enzymes is mapped between species. Hence, the number of reactions and metabolites as produced by the printModelStats function in the toolbox does not reflect the actual gene mapping between iHepatocytes2322 and each of the draft models. One way to properly examine different cod annotation outcomes, given the known fish genome duplication, is to compare the number of mapped genes from iHepatocytes2322 with each draft, with strictness s3. But the current RAVEN only generates the mapped genes included in the draft model, we thus added this feature to the getModelFromHomology function and pulled the request to RAVEN’s GitHub page (https://github.com/SysBioChalmers/RAVEN/pull/301). The Venn diagram in Fig 3 shows the output of this function. As the draft model from gadMor3 is the most complete, we used it as the baseline and included the other two as complementary sources.

**Fig 3.**
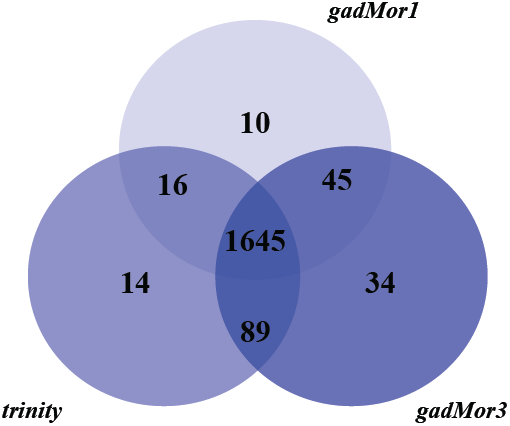
Number of genes in the Atlantic cod liver draft GEMs based on the three genome assemblies, gadMor1, gadMorTrinity, and gadMor3, when mapped to iHepatocytes2322 with strictness equal to 3.

A draft reconstruction typically contains missing genes and reactions which are gaps to be filled later. In cases where a template model is used as input, such gaps result from unmapped genes, and subsequent reactions, between the template and the target organisms. In order to examine the degree of gaps in our draft model, we considered the “Metabolism of xenobiotics by cytochrome P450” subsystem which is highly relevant to our case study. A map of this subsystem was visualized by Escher [27] as shown in Fig 4. Comparing it with the same subsystem from model iHepatocytes2322, the missing reactions are highlighted in pink. Fig 4 shows the difference before (Fig 4a) and after (Fig 4b) gap filling. The missing reactions correspond to potential gaps that need to be filled during model refinement.

**Fig 4.**
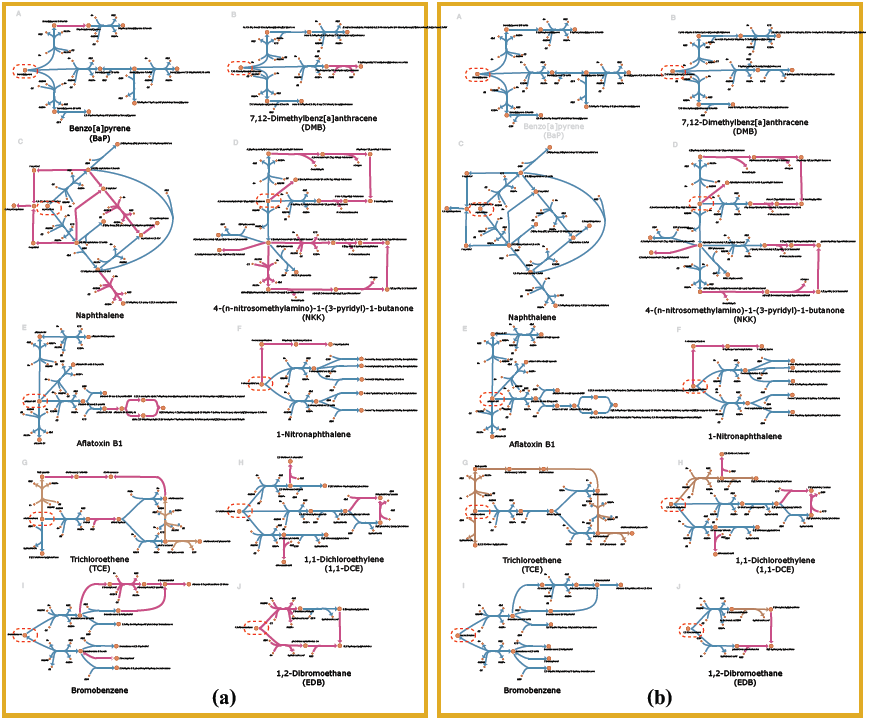
Visualization of the subsystem “Metabolism of xenobiotics by cytochrome P450” before (a) and after (b) gap filling. When enzymes are accounted for, the reactions are shown in blue, whereas the gaps are shown in pink. Dotted circles indicate the parent toxicant.

By manual curation, we set several criteria for gap filling: 1. by searching relevant literature, several reactions were identified as spontaneous reactions, i.e. not catalyzed by any enzyme, and included. 2: when the chemistry of the reaction made it clear that it belonged to a process catalyzed by a certain enzyme family (i.e. cytochrome P450 monooxygenases), they were assigned to this family even though the specific gene homolog was not identified.

## Case study of Atlantic cod exposed to benzo(a)pyrene

Mapping gene expression data to the metabolic pathway can give a better understanding of the mechanism by which a toxicant affects the organism. Here, we show a case study of *ex vivo* exposure of Atlantic cod liver slices to benzo(a)pyrene (BaP) [59]. As seen in reaction A in Figure 4b, all enzymes related to this metabolic pathway were mapped in our model following manual curation. In more details, the metabolic pathway of BaP (adapted from [47] show that there are two main biotransformation routes of BaP (Fig 5). Combining the metabolic pathway and gene expression data, we show that the main biotransformation route of BaP in Atlantic cod liver is likely towards the most reactive metabolite, benzo(a)pyrene-7,8-dihydrodiol-9,10-epoxide (Fig 5). Simultaneously, an alternate clearance pathway dependent upon the SULT and UGT enzymes is actually down-regulated, demonstrating clear preference for the primary activation pathway. This shift can be explained by the fact that BaP itself is a well-known ligand for the aryl hydrocarbon receptor (AHR), an important ligand-activated transcription factor regulating the expression of several enzymes in xenobiotic biotransformation pathways, including CYP1A, and recently characterized in cod [5]. Although this mechanism is not specific for cod and well described previously (e.g. [28, 47]), this demonstrates how a useful insight into evaluation of harmful compounds can be gained by metabolic reconstructions. The expression profiles of the genes can be found in the Supplementary materials.

**Fig 5.**
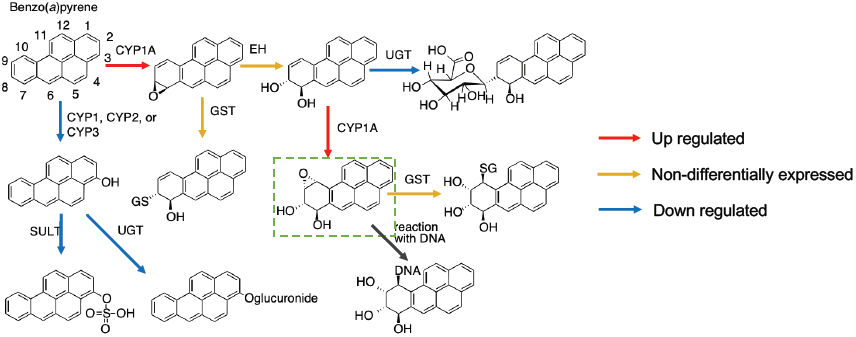
Metabolic pathways of benzo(a)pyrene (BaP) in fish based on [47] and putative changes in cod liver based on differential gene expression after exposure to BaP [59]. The upregulation of cyp1a and down-regulation of sult and ugt expression (see Supplementary materials xx) indicate that the formation of the DNA-reactive metabolite benzo(a)pyrene-7,8-dihydrodiol-9,10-epoxide is preferred.

## Discussion and future perspectives

In this paper, we presented our experience in reconstructing a draft metabolic network for Atlantic cod liver. As our genome-scale reconstruction work is still on-going, the main intention in this paper is to highlight possible challenges of working on non-model less-annotated organisms. Draft reconstruction for model species is typically the fastest phase in the process, due to the abundance of biochemical knowledge and annotations. However, it becomes less straightforward with a lack of such data. Thanks to enabling computational tools, an alternative is to use existing generic or tissue-specific GEMs that fall within the context of the target reconstruction. In our case, a human liver model was used as a template to generate a draft cod liver model. The fact that it is a well-annotated consensus liver GEM compensated for known physiological and metabolic differences between cod and human, which could be resolved in the next model curation phases.

As demonstrated for the “Metabolism of xenobiotics by cytochrome P450” subsystem, gap filling is an essential part of model curation that need to be applied to all the included subsystems. We survey main gap filling approaches and then describe subsequence reconstruction steps. Several tools were presented to help automate this step. Among those approaches is Meneco [43], a topological gap filling approach, written in Python, that identifies missing reactions through reformulating the gap filling problem as a combinatorial optimization problem. fastGapFill [51] is available in the COBRA toolbox. It selects the set of reactions that optimize a linear score representing the quantitative metabolic production of the metabolic network. Other existing tools can identify missing reactions by integrating taxonomic [7], compartment modularity [38], or phenotypic knowledge such as experimental flux data [12, 23, 30, 55]. However, approaches relying on complementary knowledge to perform the gap filling process are not applicable to less-annotated organisms. Accordingly, fastGapFill from the COBRA toolbox could be a suitable choice to automatically fill the gaps in the draft model. Following this step, the refined set of reactions needs to be manually curated since some of them might be added arbitrarily to fulfill selected model criteria, such as model connectivity, by the gap filling tool. The model is then ready for evaluation using flux balance analysis (FBA) [41] and phenotype datasets. FBA aims to simulate the cell behavior through finding a set of optimal metabolic fluxes which optimizes the biomass production (i.e. cellular growth). The simulation results from FBA are compared with experimental values of biomass growth and metabolic fluxes and the model is subjected to refinement until it behaves as desired. The latter allows the reconstructed metabolic network to be analyzed using constraint-based modeling approach. Once those steps are completed, the model can be used to simulate different biological conditions and study the metabolic potential of the organism. For example, it is possible to perform gene knockout studies in silico to identify essential genes for various biological conditions using the constraint-based model of the reconstructed network. Thus, insights can be gained into how important physiological processes in the organism, including energy metabolism, growth and reproductive processes, can be affected by environmental toxicants.

## Acknowledgments

This work was supported by the Research Council of Norway to the Centre for Digital Life Norway (DLN) project dCod 1.0: decoding the systems toxicology of Atlantic cod [grant number 248840] and the Norwegian research school in bioinformatics, biostatistics and systems biology (NORBIS).

